# Theta sweeps accelerate cognitive map learning by broadcasting information to unvisited regions

**DOI:** 10.64898/2026.09.26.754738

**Authors:** Tianhao Chu, Jinfan He, Zilong Ji, Fang Fang, Si Wu

## Abstract

To support flexible behavior, animals need to learn map-like representations of environments from limited experiences, but the underlying neural mechanisms remain unclear. Here, we propose that theta sweeps, a prominent feature in the activities of grid and place cells during active movement, can broadcast acquired sensory information to swept unvisited regions, accelerating cognitive-map learning. To test this idea, we built a network of direction, grid, and place-cell modules, in which theta modulation and firing-rate adaptation generate theta sweeps, while Hebbian plasticity enables place cells to bind environmental observations. Simulations and ablations showed that the dorsal–ventral grid organization produces multiscale, independent sweeps broadcasting acquired sensory information to a broad, fan-shaped unvisited area, enabling unvisited regions to build place-environment associations. The model further suggests that declining theta power as the environment becomes familiar acts as an annealing process, shifting from learning general information about the environment to making local refinements. The model makes three testable predictions: strong theta modulation accelerates early learning, environments with broader spatial correlations favor longer theta sweeps, and theta sweeps progressively shorten as environments become familiar. Together, our results support theta sweeps as a mechanism for accelerating cognitive-map learning from limited explorations, and it sheds light on developing brain-inspired algorithms for building world models.

## 1 Introduction

Cognitive map theory proposes that the brain builds an internal mental representation of the physical environment and abstract concepts to support memory, navigation, and decision-making (Whittington et al., 2020). By separating sensory content from relational structure, previous cognitive-map models accelerate learning across environments with shared structure: only new observation bindings need to be learned (Whittington et al., 2020; Chandra et al., 2025; George et al., 2021). In these models, the extent of map learning is assessed by how accurately the model predicts sensory observations (Whittington et al., 2020), an approach also used to evaluate learned world models (Ha & Schmidhuber, 2018). In graph terms, this shifts learning the edges that encode relations (Figure 1a) to learning how sensory observations are bound to the shared structure by the node (Figure 1b). However, node-level learning remains inefficient, especially in continuous environments: learning sensory content across the map still requires dense spatial coverage, which is infeasible and inefficient (Figure 1c). It is unclear how neural circuits use sparse experience to improve sensory predictions at unvisited locations.

**Figure 1.**
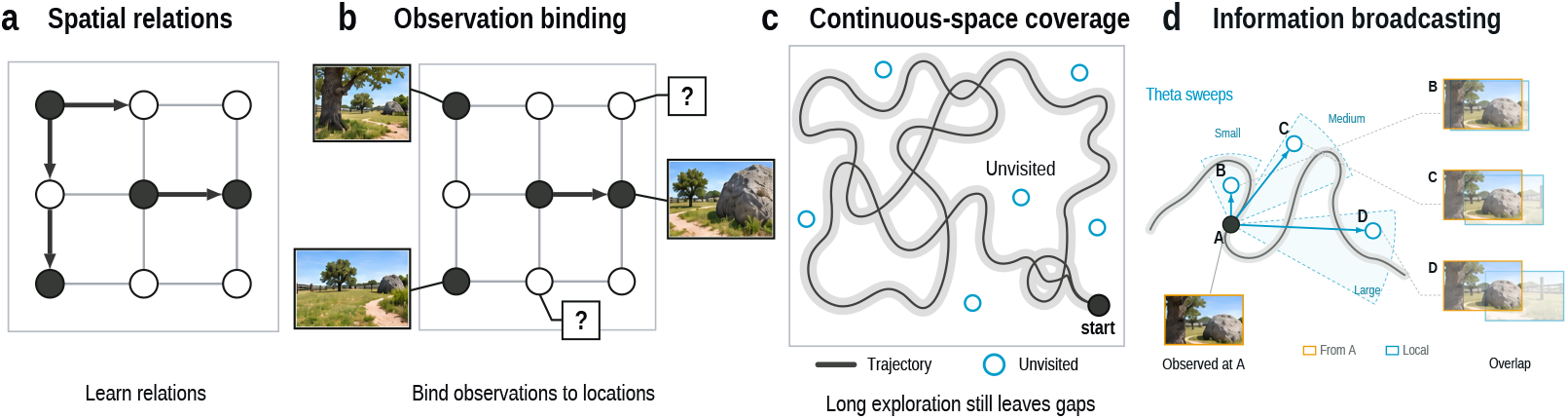
Spatial learning and information broadcasting. Schematics of (a) relational structure, (b) observation binding, (c) incomplete physical coverage, and (d) broadcasting an observation from A to internally sampled B–D. Orange and blue frames show source and local sensory content, respectively, using equal-width crops of one scene.

Theta sweeps may provide a promising neural mechanism for structured sampling of unvisited locations. Recent studies (Vollan et al., 2025) have shown that, when animals freely explore an environment, locations decoded from grid-cell population activity sweep from the animal’s current position toward either side of its heading direction across successive theta cycles (roughly 10 cycles per second), alternating between left and right. Across grid-cell modules, these sweeps vary with spatial scale and collectively form a fan-shaped sampling pattern around the animal (Ji et al., 2026). We therefore hypothesize that this theta-paced sampling accelerates cognitive-map learning through *information broadcasting*. While the animal is at A, sweeps activate grid cells coding for locations B, C, and D, recruiting hippocampal place-cell representations that synaptic plasticity associates with the sensory observation at A (Figure 1d). Spatial correlations in sensory content make this observation informative about these unvisited locations. Over successive sweeps, each location’s representation integrates observations from multiple visited locations, allowing its sensory content to be estimated without visiting it directly.

To test this hypothesis, we developed an entorhinal–hippocampal network that couples theta-sweep dynamics (Ji et al., 2025) to plastic sensory associations, allowing us to examine how internally sampled locations shape cognitive-map learning. Our model (Figure 2a) includes multiple grid modules with different spacings, reflecting the modular organization and dorsal–ventral scale variation in the medial entorhinal cortex (MEC) (Stensola et al., 2012). These modules project to hippocampal (HPC) place-cell populations with different field sizes through Gaussian-weighted connections, reflecting the dorsal–ventral increase in place-field extent (Kjelstrup et al., 2008). We use Hebbian plasticity to bind HPC activity to sensory activity modeled in lateral entorhinal cortex (LEC), where object-related responses have been recorded (Deshmukh & Knierim, 2011), as in other cognitive-map models (Chandra et al., 2025). These HPC-to-LEC connections learn online during exploration, binding current observations to active hippocampal representations; all other connections remain fixed.

**Figure 2.**
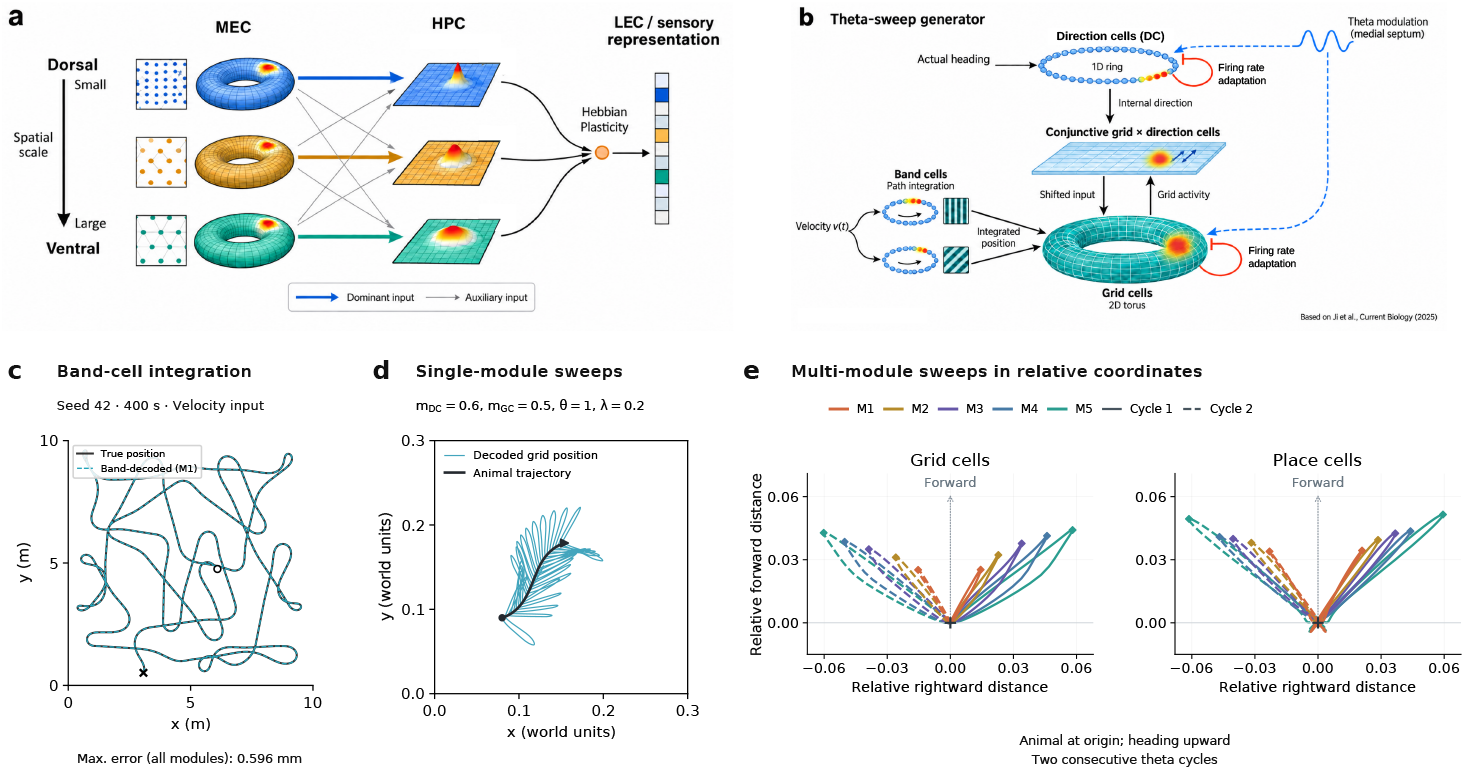
Circuit organization and trajectory measurements. (a) MEC–HPC modules and sensory readout. The illustration shows three module pairs; simulations use five. (b) Sweep generation and band-cell reference pathway. Red feedback loops indicate firing rate adaptation in direction cells and grid cells. (c) Physical (dark) and decoded position (dashed cyan) from band cell of a 400 s velocity integration. (d) Physical (dark) and theta sweeps (blue) from one grid module. (e) Theta sweep trajectories from grid-(left) and place cells (right) relative to the agent, with heading upward; colors representing different modules. Numerical panels show individual runs.

We show theta sweeps accelerate sensory-map learning and improve predictions at unvisited positions relative to both the no-sweep network and the Vector-HaSH control. Ablations and parameter sweeps clarify how the circuit’s biological organization contributes to learning. Dorsal–ventral grid and place modules generate sweeps with different directions and lengths, covering more space in less time and accelerating learning. Hippocampal recurrence, motivated by recurrent connectivity in CA3 (Guzman et al., 2016), improves learning by supporting localized representations for binding. We also find a tradeoff between early learning speed and final prediction accuracy: strong broadcasting speeds initial learning, whereas weaker broadcasting improves precision as exploration fills the map. An annealing schedule that progressively reduces theta modulation combines rapid initial learning with lower final prediction error than either fixed-modulation condition. This suggests that reduced hippocampal theta power in familiar environments (Penley et al., 2013) may serve an annealing function, allowing animals to learn environmental information rapidly in the early stage of exploration and improve prediction accuracy as experience accumulates.

The model makes three testable predictions. First, stronger theta modulation should accelerate learning during early exploration. Second, the theta-sweep length that best supports learning should depend on the spatial correlation length of LEC population activity, with broader correlations favoring longer sweeps across animals and sensory environments. Third, theta sweeps should progressively shorten as familiarity increases and theta modulation declines, as predicted by the annealing account. Together, these results support theta sweeps as a mechanism for rapid cognitive-map learning from sparse experience and suggest information broadcasting as a potential strategy for world models to rapidly learn spatial representations of unfamiliar environments.

## 2 Related work

### Accelerating cognitive-map learning from edges to nodes

Models such as the Tolman– Eichenbaum machine and Vector-HaSH support cognitive-map learning by separating sensory content from spatial structure (Whittington et al., 2020; Chandra et al., 2025). These models learn how sensory information is bound to locations on the reusable structure rather than rebuilding the cognitive map from scratch. We build on this framework by using theta sweeps to extend sensory binding beyond visited locations, and compare against a Vector-HaSH-based control to test whether this spatial sampling accelerates learning.

### Neural circuit mechanisms for theta sweeps

Our spatial sampling circuit builds on models of theta-sweep generation and band-cell path integration. Adaptation-based attractor models generate place-cell theta sweeps (Chu et al., 2024), while coupled direction and grid-cell networks generate left–right-alternating sweeps (Ji et al., 2025). We adopt this theta-sweep generator to sample positions beyond the animal’s direct experience. Periodic band representations encode displacement along preferred directions and drive grid-cell networks (Bush & Burgess, 2014; Chu et al., 2025), providing the basis for the position reference that anchors theta sweeps. Together, these models provide the spatial dynamics on which we build hippocampal-to-sensory binding, allowing us to test how internally sampled positions contribute to learning sensory predictions.

### Computational functions of theta sweeps

Several functions have been proposed for theta sweeps, including planning, accelerated learning, and memory retrieval. Computational models have formalized these hypotheses through trajectory sampling (Ujfalussy & Orbán, 2022), sequence recall (Jensen & Lisman, 1996), and reward-based credit assignment (George, 2023). For accelerated learning, theta phase precession was proposed to compress activity sequences into the STDP window, facilitating learning of successor-like representations in CA3-to-CA1 connections (George et al., 2023). These representations predict future occupancy rather than sensory content. Left–right sweeps accelerate learning of recurrent place-cell connections by extending neural activity beyond the physical trajectory (Marshall et al., 2025). We extend this principle of expanding the spatial extent of learning by asking how information gathered during exploration can be broadcast to unvisited areas to improve sensory predictions.

## 3 An MEC–HPC neural circuit model for learning with theta sweeps

### 3.1 Model architecture

Our model comprises grid-cell modules in MEC, place-cell networks in HPC, and a sensory population in LEC (Figure 2a). MEC serves as a theta-sweep generator, whose circuit and dynamics are described in Section 3.2; its output drives sweeps in downstream HPC networks. This organization is motivated by recent findings showing that hippocampal sweeps inherit entorhinal sweeps (Vollan et al., 2025).

We organize MEC and HPC into multiple grid–place module pairs to support precise position coding across scales and broad spatial sampling through sweeps in different directions. This organization is consistent with the dorsoventral increase in grid spacing and place-field size (Brun et al., 2008; Stensola et al., 2012; Kjelstrup et al., 2008). Motivated by the scale-dependent sweep directions observed across MEC modules (Ji et al., 2026), we allow different grid modules and their downstream HPC populations to sweep along different directions, collectively sampling a fan-shaped region around the animal. See Appendix A.2 for spatial scales and module settings.

Each place-cell network combines inputs from multiple grid modules to resolve the spatial ambiguity of periodic grid codes (Solstad et al., 2006). Input is strongest from the corresponding grid module located proportionally on the dorsoventral axis and weaker from the remaining modules, consistent with entorhinal–hippocampal projection topography and previous models (Dolorfo & Amaral, 1998; Lyttle et al., 2013) (see Appendix A.2 for the projection fields and input weights).

We model *M* place-cell populations, indexed by *k* = 1, *…, M*, with cells in each population encoding different locations of the environment. Within each population, place cells have the same field size *a*_*p,k*_, while field sizes differ across populations. Each population forms a recurrent neural network (Guzman et al., 2016) with local excitation and global inhibition (Wu et al., 2008), modeling the recurrent connections found in the hippocampal CA3 region. For simplicity, we restrict recurrent connections within the same population, with excitatory weights

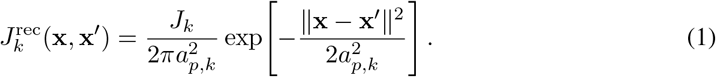

Here **x** and **x**^*′*^ are place-field centers within population *k*, and *J*_*k*_ denotes the connection strength. We model the dorsoventral correspondence between grid-cell modules and place-cell populations by connecting their spatial scales through *a*_*p,k*_ = *λ*_*k*_/(2*π*), where *λ*_*k*_ is the spacing of grid cells in the *k*th grid module (see Appendix A.2 for implementation details).

We add global inhibition to presynaptic input *u*_*k,i*_, so that firing rate *r*_*k,i*_ is normalized:

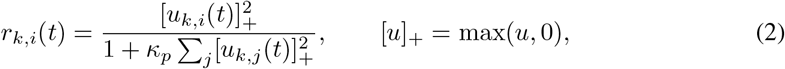

where *κ*_*p*_ represents the inhibition strength. Together with local excitation, this inhibition supports a localized activity bump. See Appendix A.1 for place-cell dynamics.

In the model, LEC neurons are modeled as landmark vector cells (LVCs), each tuned to a preferred direction and distance relative to a landmark. Together, these cells represent landmark-relative position in polar coordinates, with activity driven by sensory input at the animal’s current location (see Figure 7e,f). Synaptic connections between LVCs and place cells are updated through a delta-Hebbian rule (Widrow & Hoff, 1960):

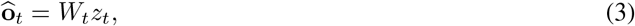

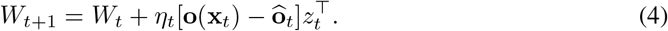

Here *η*_*t*_ is the learning rate, and *z*_*t*_ = *f* (*r*_*t*_) maps the HPC firing-rate vector *r*_*t*_ to the activity supplied to the sensory readout. Delta-Hebbian plasticity continually updates the synaptic weights *W*_*t*_ to reduce the prediction error between hippocampal output and current sensory input. During a theta sweep, HPC place cells representing locations around the animal are sequentially activated, while LEC continues to represent sensory input at the animal’s current location. This plasticity associates the current sensory input with locations represented along the sweep, allowing a single observation to update predictions even at unvisited locations. When sensory inputs vary smoothly across space, these updates support learning at unvisited locations and reduce the need to sample each location directly. See Appendix A.3 for sensory tuning, the definition of *f*, and synaptic learning details.

### 3.2 Theta-sweep generator

To generate theta sweeps in grid modules, we construct a direction-cell (DC) circuit that drives grid cells through an intermediate population of conjunctive grid-by-direction cells (Figure 2b), following Ji et al. (2025). With theta modulation and firing-rate adaptation, the direction population generates alternating left–right sweeps of represented direction, which drive corresponding theta sweeps of represented location in grid cells (Figure 2d). A positional input anchors each theta sweep to the animal’s current position. We model this positional input using the upstream path integrator of Chu et al. (2025). Two band-cell populations integrate velocity along axes separated by 60^*°*^, matching the hexagonal lattice geometry of grid-cell firing fields. Their combined activity provides the positional input to the downstream grid-cell network (see Appendices A.1 and A.4).

We first reproduce the basic behavior of the two component models. For band-cell path integration, the network starts from a known initial position and receives velocity input for 400 s. Position decoded from band-cell activity closely follows the true trajectory, with a maximum accumulated error of 0.596 mm across modules (Figure 2c). For theta-sweep generation, Figure 2d shows a single grid-cell module as the animal follows a curved path. The decoded location repeatedly moves away from and returns toward the animal’s current position, sweeping ahead and alternating between left and right of the heading direction (Figure 2d).

We then examine these sweeps in our full MEC–HPC–LEC model (Figure 2a), with multiple grid modules projecting to place-cell populations at different spatial scales. Grid-module sweep angles increase along the MEC dorsoventral axis, alongside a corresponding increase in directional sweep angles in the upstream parasubiculum (Ji et al., 2026). Motivated by this organization and its proposed adaptation-based mechanism, we couple each grid module to a separate direction-cell module, with stronger adaptation in the DC modules driving larger-scale grid modules. Stronger adaptation produces wider left–right angular sweeps in DC activity, which in turn steer grid-cell sweeps along different directions across modules (see Appendix A.1 for implementation details). Decoded grid-cell and place-cell trajectories are shown over two consecutive theta cycles in an animal-centered reference frame, with the animal’s heading aligned upward (Figure 2e). Grid-cell sweeps extend to opposite sides of the heading in successive cycles, and the downstream HPC populations inherit this alternation from their grid-cell inputs. Larger-scale modules show longer sweeps in both grid-cell and place-cell populations.

## 4 Results

### 4.1 Theta sweeps accelerate learning and improve predictions at unvisited locations

We tested whether theta sweeps accelerate learning by using sensory observations at visited locations to improve predictions at unvisited locations. The simulated animal explored a 10 × 10 m arena containing four landmarks (Figure 3a). For the main-text results, we modeled LEC neurons as landmark-vector cells (LVCs), which fire when the animal is at a preferred distance and direction from a landmark (Figure 3b). During exploration, plastic HPC–LEC synapses associated place-cell activity with LVC activity; all other connections remained fixed. To test whether the learning benefit depended on LVC encoding, we also compared learning with and without sweeps using object-specific-vector and boundary-distance representations. Both alternatives yielded the same qualitative benefit (Appendix C.2).

**Figure 3.**
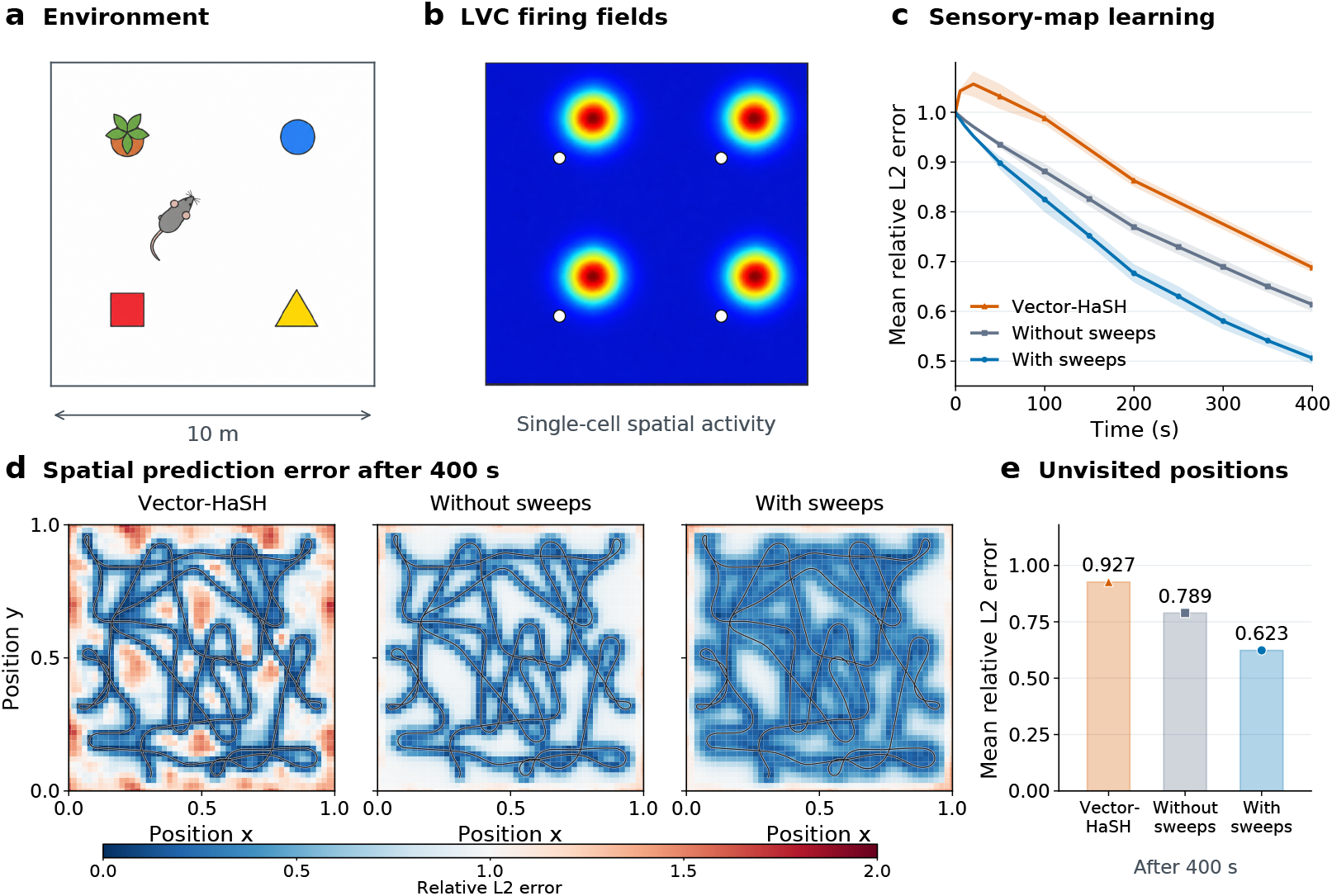
Theta sweeps accelerate learning of a map of the environment and improve predictions at unvisited positions. (a) Schematic arena containing four landmarks. (b) Schematic spatial firing fields of a single LVC, at the same direction and distance from each landmark (white circles). (c) All-position relative-*L*_2_ error for the Vector-HaSH model, and our no-sweep-, and sweep-based models under the same exploration trajectory (mean ± SD over three trajectories). (d) Spatial heatmaps of relative-*L*_2_ error. At each query location, error is the *L*_2_ norm of the difference between predicted and target LEC activity vectors, normalized by the target vector’s *L*_2_ norm (Equation 25). Maps share a color scale; black lines show the physical trajectory. (e) Unvisited-position errors among three models.

Theta sweeps accelerated sensory-map learning, reducing the mean relative *L*_2_ error in predicted LEC activity across the arena faster than either the no-sweep model or Vector-HaSH (Figure 3c). This advantage is consistent with sweeps extending each sensory learning update to nearby, internally represented locations, allowing the same exploration to improve predictions over a broader area. We chose Vector-HaSH as a baseline because it also binds sensory observations to a grid-based hippocampal spatial representation, providing a comparison with an established map-learning model that lacks theta sweeps. All three models had matched numbers of grid, hippocampal, and sensory neurons and experienced the same trajectories and sensory inputs for 400 s. Final error was 0.506 ± 0.012 with sweeps, compared with 0.614 ± 0.014 without sweeps and 0.688 ± 0.010 for Vector-HaSH (mean ± SD across three trajectories), corresponding to reductions of 17.5% and 26.4%, respectively.

Even without sweeps, our model learned faster than Vector-HaSH, reaching a 10.8% lower final error. Two circuit features distinguish it from the Vector-HaSH control: recurrent connections within each CA3 population and spatially structured grid-to-place projections, which replace the random projections in the control. These connections favor a single localized activity bump within each HPC population and similar firing patterns at nearby locations (see the recurrence ablation in Section 4.2). This spatial organization allows sensory experience to generalize across neighboring place representations.

We compared the three models after 400 s of matched exploration at the same unvisited query locations, defined as positions more than 0.15 m from every point on the physical trajectory. With learning disabled, we computed each model’s average relative *L*_2_ error between predicted and target LEC activity at these locations. The spatial heatmaps in Figure 3d show that our model with theta sweeps predicted sensory activity more accurately in the gaps between explored paths, with low-error regions extending farther from the animal’s trajectory. Figure 3e quantifies this difference: mean error at unvisited locations was 0.623 with sweeps, compared with 0.789 without sweeps and 0.927 for Vector-HaSH in the illustrated trajectory, reductions of 21.0% and 32.7%, respectively. Improved hippocampal predictions outside the explored path therefore explain why theta sweeps accelerated learning of the overall sensory map. Evaluation details and separate regional learning curves are given in Appendices B.3 and C.2.

### 4.2 Multiple place-cell scales and CA3 recurrence support sensory-map learning

We tested how place-cell populations with different field sizes, driven by grid-cell modules with corresponding spatial scales, and recurrent excitation within CA3 contribute to learning a map of sensory predictions across the environment.

We used only place-cell population M1, paired with the smallest grid spacing (Table 2), for output normalization and sensory readout. This ablation slowed learning and increased prediction error (Figure 4a). With a single module, place-cell activity covered the environment more slowly during the same physical exploration (Figure 4b). Sweeps at multiple spatial scales recruit place cells representing a broader region, allowing the current sensory experience to be associated with more locations (see Appendix B.3 for the coverage measure).

**Figure 4.**
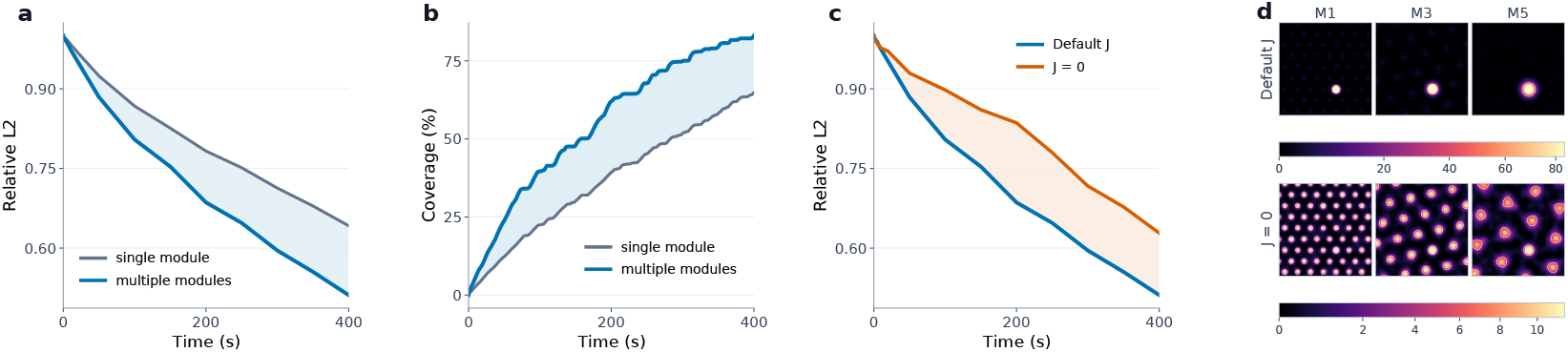
Multiple spatial scales and CA3 recurrence support sensory learning. (a,b) Learning from a single module (gray) or multiple modules (blue), and the cumulative spatial coverage of their decoded place activity. (c,d) Learning and place-cell activity snapshots with and without recurrent excitation (*J* = 0); snapshot rows use separate color scales. All panels use one matched 400 s exploration trajectory.

Removing recurrent excitation within CA3 (*J* = 0) also slowed learning and increased prediction error (Figure 4c). Recurrence shapes place-cell activity in two ways. First, local recurrent excitation favors a single localized activity bump, whereas without recurrence, periodic grid input produces multiple spatially separated peaks (Figure 4d). Second, recurrence can support spatial continuity: activity from the preceding moment favors nearby place cells over distant competing peaks. In a two-module circuit, removing recurrence caused the decoded bump position to jump between distant locations, whereas recurrent connections maintained continuous movement under the same grid input (see Appendix C.3 and Figure 8).

#### 4.3 Theta power annealing balances learning speed and prediction accuracy

Information broadcasting introduces a potential speed–accuracy tradeoff in sensory-map learning: associating an observation with a broad range of place representations can accelerate learning from sparse experience, but sensory differences between locations can limit the precision of these associations. In the model, theta modulation rhythmically weakens the inputs anchoring direction-cell and grid-cell activity to the animal’s heading and position, allowing firing-rate adaptation to drive sweeps away from these references. We tested this tradeoff by extending the same exploration trajectory to 4,000 s and comparing fixed strong theta modulation with zero modulation, which maintains constant anchoring and largely suppresses theta sweeps. Strong modulation reduced prediction error faster early in learning, but its advantage diminished as exploration covered more of the environment (Figure 5a). Later in exploration, zero modulation achieved lower all-position error (Figure 5a). This crossover did not reflect a loss of the broadcasting benefit at unvisited locations: strong modulation still produced lower error there than zero modulation (Appendix C.2, Figure 7d). Later in learning, however, this generalization benefit was outweighed by the greater prediction accuracy at visited locations under zero modulation.

**Figure 5.**
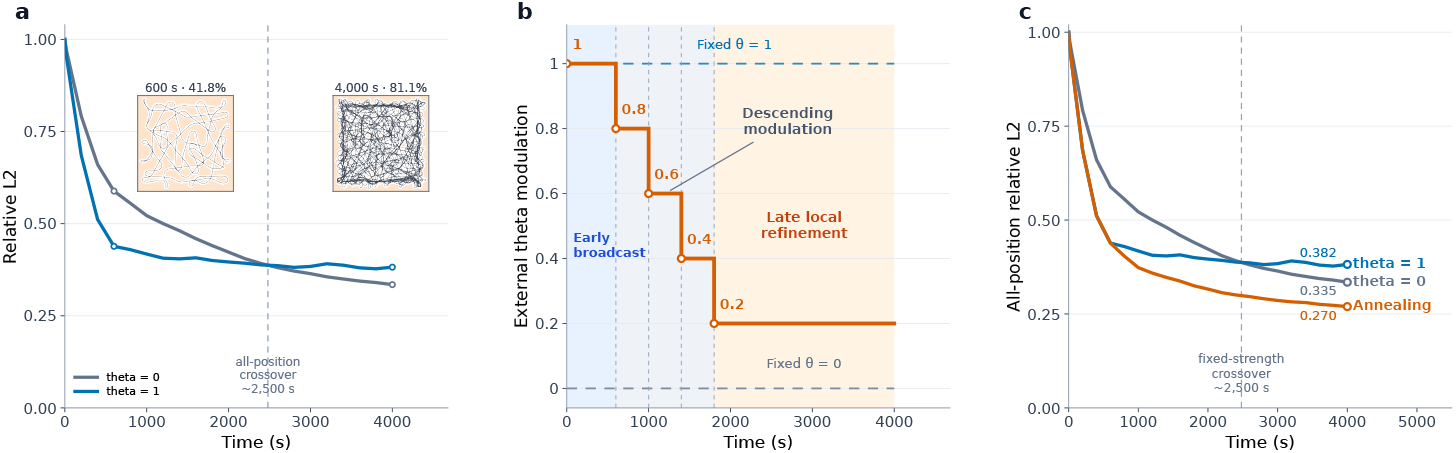
Theta annealing balances learning speed and prediction accuracy. (a) Learning under fixed modulation; insets show physical coverage at 600 and 4,000 s, with unvisited regions in peach. (b) Theta-annealing schedule. (c) Fixed-strength and annealed learning curves. All conditions use the same 4,000 s trajectory and fixed learning coefficient.

To combine these benefits, we introduced a theta-annealing schedule that progressively reduces modulation as exploration proceeds (Figure 5b; see Appendix B.6 for the schedule). Lower modulation keeps neural activity more closely anchored to the animal’s location, reducing sweep amplitude and the spatial reach of information broadcasting. As direct observations accumulate, this shifts learning from broad spatial generalization toward local refinement. Annealing retained the early learning advantage of strong modulation and reached lower final all-position error than either fixed condition (Figure 5c). This annealing effect aligns with the lower hippocampal LFP theta power observed in familiar than novel environments (Penley et al., 2013). Our results show why reduced theta power might be useful during spatial navigation: a decline in theta power as the environment becomes familiar may serve an annealing function, balancing learning speed and prediction accuracy through reduced information broadcasting, and finally reducing the all-position relative *L*_2_ error.

#### 4.4 Model predictions from theta modulation and adaptation

Our model makes predictions that can be tested with biological data. We focused on theta-modulation strength and firing-rate adaptation, two key components of theta-sweep generation.

First, we predicted that stronger theta modulation should accelerate learning during early exploration. We systematically varied theta-modulation strengths (see Appendix B.1 for details). Stronger modulation produced a faster decline in prediction error (Figure 6a).

**Figure 6.**
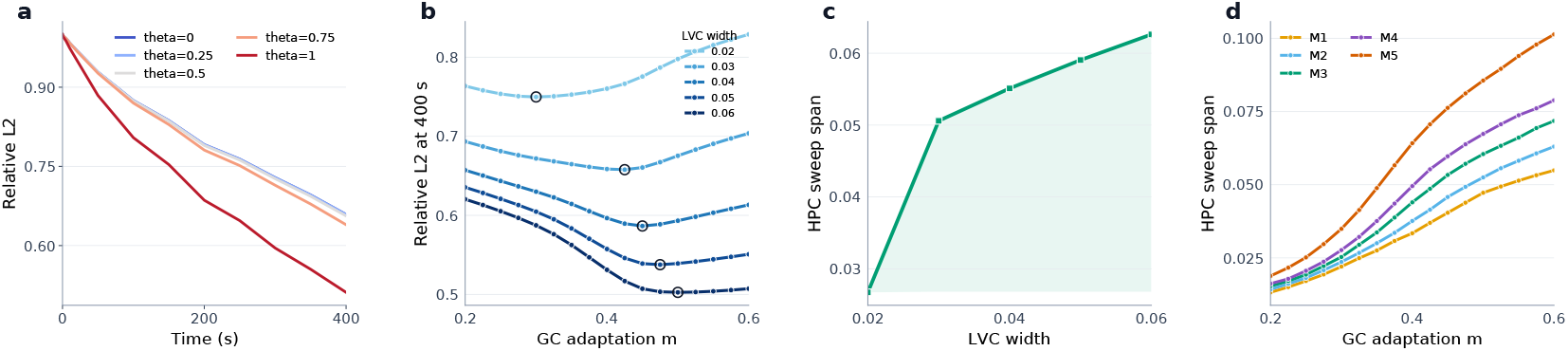
Predictions from theta modulation and adaptation. (a) Learning under different theta-modulation strengths on one matched trajectory. (b) Grid-adaptation scan across LVC widths; open rings mark the lowest validation error for each width. (c) Mean HPC sweep span at the adaptation selected for each LVC width. (d) HPC sweep span in each population as grid-cell adaptation increases. Widths and spans use model length units; (b–d) show means over three validation trajectories. All learning runs last 400 s.

**Figure 7.**
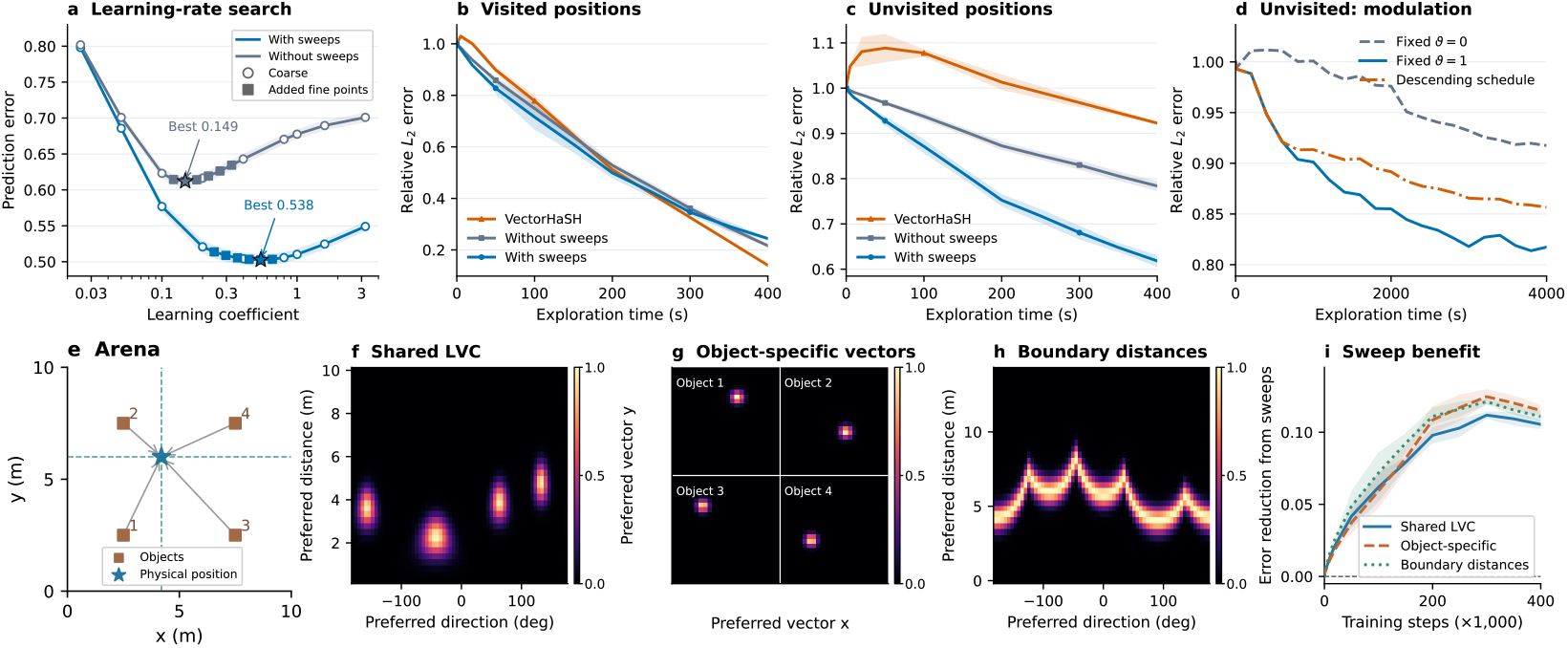
Learning-coefficient selection, spatial generalization, and sensory encodings. (a) Coarse (circles), fine (squares), and selected (stars) coefficients; validation error at 400 s (mean ± SEM). (b,c) Visited and unvisited errors (mean ± SD), with regions fixed by each complete 400 s trajectory. (d) Unvisited error for regions fixed by the complete 4,000 s trajectory. (e) Scene and query position. (f–h) Shared landmark-vector, object-specific-vector, and boundary-distance responses at that position, shown relative to each encoding’s amplitude. (i) Reduction in all-position error, Δ*E* = *E*_without_ − *E*_with_ (mean ± SD). Panels a–c,i summarize three trajectories; d shows one trajectory.

Second, we predicted that the theta-sweep length that best supports learning should depend on the spatial correlation length of the sensory representation, with broader correlations favoring longer sweeps. We tested this by scanning grid-cell adaptation, which controls sweep range: stronger adaptation increased HPC sweep span across populations (Figure 6d), and prediction error after a fixed period of exploration varied nonmonotonically with adaptation, first decreasing and then increasing (Figure 6b). Short sweeps limit spatial generalization, whereas excessively long sweeps can associate an observation with locations whose sensory content differs, producing an interior optimum. Consistent with the prediction, broader LVC tuning fields favored stronger adaptation at this optimum, and the associated HPC sweep span increased with LVC field width (Figure 6c). Broader LVC fields preserve greater sensory overlap at a given sweep distance, allowing observations to improve predictions farther away. See Appendix B.7 for parameter settings and selection, and Appendix B.3 for sweep measurements.

Third, we predicted that theta sweeps should progressively shorten as the environment becomes familiar and theta modulation declines; the annealing simulation in Section 4.3 establishes the computational basis of this prediction.

### 5 Conclusion and Discussion

Theta sweeps have been widely observed in the activities of grid and place cells when animals are in active motion. Previous studies have suggested that theta sweeps have the functions of forward planning, enabling sequence learning by generating phase precession, and memory retrieval (Ujfalussy & Orbán, 2022; George et al., 2023; Jensen & Lisman, 1996). Here, we proposed that theta sweeps contribute to broadcasting sensory observation to nearby unvisited regions, utilizing the statistical property that nearby environmental information is highly correlated. In this way, the neural system accelerates the learning of a cognitive map of the whole environment through limited explorations. We built a computational model composed of direction, grid, and place-cell modules, in which theta modulation and firing-rate adaptation generate theta sweeps, while Hebbian plasticity enables place cells to bind environmental observations. We showed that the dorsal–ventral grid organization produces multiscale, independent sweeps, which broadcast acquired sensory information to a broad, fan-shaped unvisited area, enabling the fast construction of the cognitive map in unvisited regions.

#### Experimental predictions

Our model makes three experimentally testable predictions. First, stronger theta modulation should accelerate learning during early exploration. Second, in the framework of information broadcasting, the range of theta sweeps correlates with the spatial correlation of sensory observations: unfamiliar environments with broader spatial correlations favor longer theta sweeps in the early stage of learning. Third, within the same environment, the range of theta sweeps undergoes an annealing process as exploration proceeds. The annealing prediction is consistent with the experimental finding that hippocampal LFP theta power in familiar environments is lower than that in novel environments (Penley et al., 2013). Further neuroscience experiments on behaving animals are needed to test our model predictions.

#### Limitation and future direction

A limitation in the current model is that it assumes predefined LEC population activity with hand-designed tuning functions. This simplification allows us to control spatial correlations and isolate the broadcasting mechanism, but it does not reflect the diversity and complexity of sensory observations in reality. Our future work is to extend the current framework to include place-sensory binding of high-dimensional real images in natural environments. This needs to resolve how information broadcasting works on sensory representations learned from experiences, as spatial correlations of the latter no longer follow the prescribed tuning functions used in the present study. This extension will help us to develop brain-inspired algorithms for building cognitive maps (world models) of unfamiliar environments from limited explorations.

## AI use statement

Generative AI assisted in organizing the authors’ model description, drafting manuscript text, preparing the LaTeX project, and generating the conceptual artwork in Figure 1 and Figures 2(a,b) and 3(a,b). Figures 2(c–e), 3(c–e), 4–6, and the numerical panels of Figure 8 were plotted from saved neural simulations and learning results. The continuity schematic in Figure 8a was adapted from the authors’ drawing.

**Figure 8.**
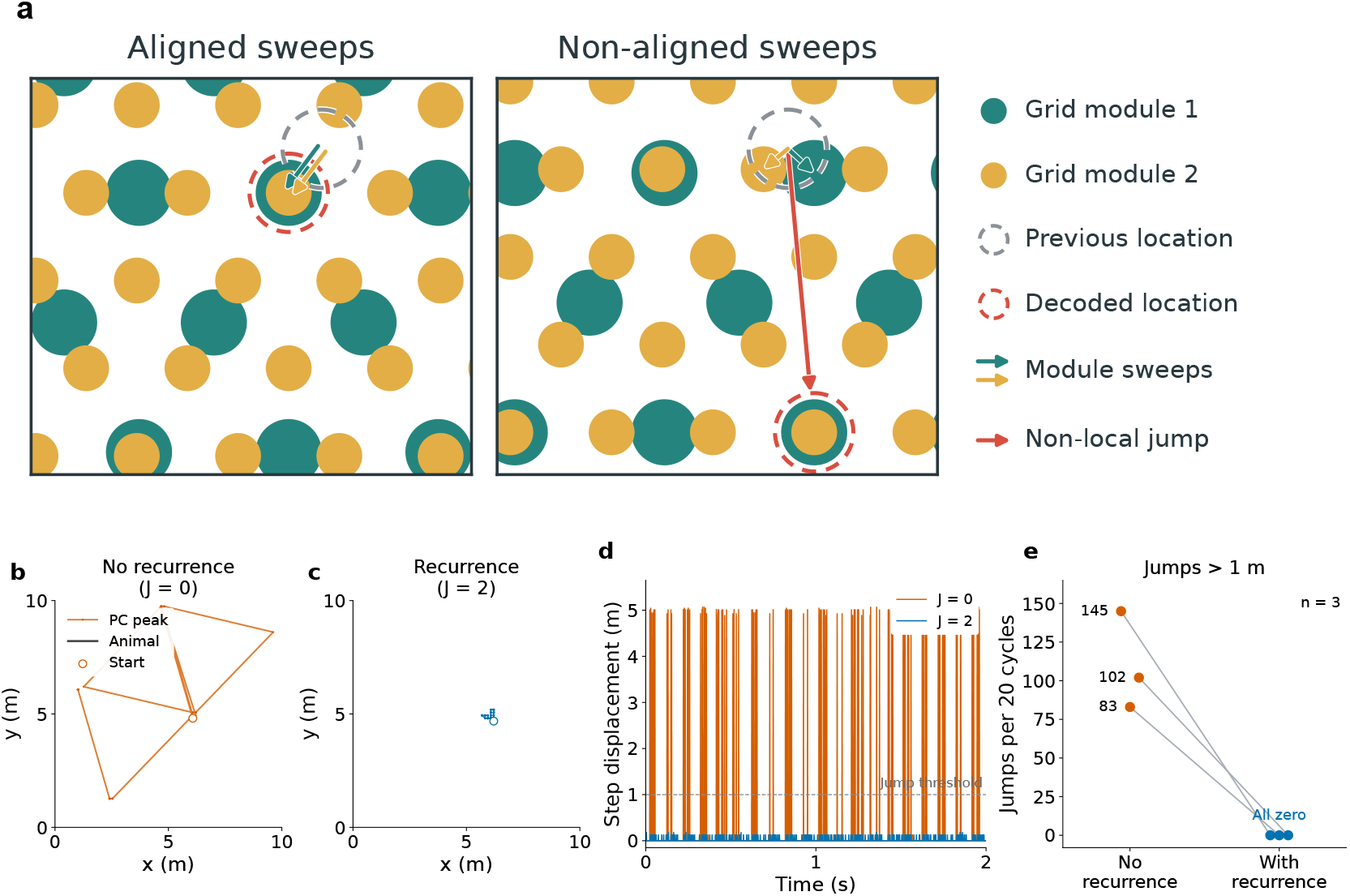
Recurrent excitation prevents non-local peak switches in a two-module circuit. (a) Aligned sweeps preserve the nearby overlap of two periodic representations, whereas non-aligned sweeps can favor a distant overlap. (b,c) Maximum-rate PC decoding over the first two recorded theta cycles of the same trajectory, without and with recurrence; both use identical 0–10 m coordinate ranges and equal axis scaling. Black lines show the animal’s path and open circles mark the initial decoded position. (d) Within-cycle displacements for this example; the dashed line marks the non-local-jump threshold. (e) Jump counts for the three paired trajectories. See Appendix B.5 for the experimental settings and jump definition.

## Reproducibility statement

Section 4.1 defines the task and evaluation metric. For model equations, units, and parameter details, see Appendix A; for experiment and evaluation details, Appendix B; and for additional experimental results, Appendix C.

## A Model architecture and settings

### A.1 Dynamics of DC, MEC, and HPC populations

The circuit contains multiple DC rings, MEC grid sheets, and HPC place-cell populations. Each DC ring supplies directional input to one MEC module, whose activity then projects to HPC populations at different spatial scales. Unless otherwise specified, the circuit has *M* = 5 modules of each type. Times are expressed in seconds and angles in radians unless specified otherwise. Activity and input amplitudes use arbitrary model units (a.u.); connection and inhibition coefficients retain the normalization of the model equations.

DC, MEC, and HPC populations use adaptation-based attractor dynamics (Chu et al., 2024).

#### Shared activity dynamics

For cell *a* in a population, synaptic activity *u*_*a*_ and adaptation *v*_*a*_ evolve as

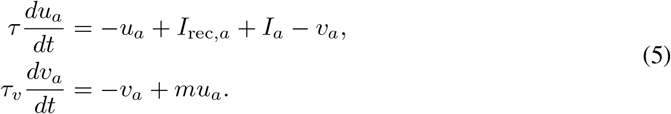

Here *t* is time, *I*_rec,*a*_ is recurrent input, and *I*_*a*_ is input from outside the population. The time constants *τ* and *τ*_*v*_ govern activity and adaptation, and *m* sets adaptation strength. Adaptation provides delayed negative feedback to active cells, allowing the activity bump to move. This mechanism is motivated by MEC adaptation measurements (Yoshida et al., 2013) and models of bump motion (Mi et al., 2014).

Firing rates follow the squared-activity normalization in Equation 2, with inhibition coefficients *k*_*h*_, *k*_*g*_, and *κ*_*p*_ for DC, MEC, and HPC, respectively. Each population has its own normalization pool. Table 1 gives the population parameters. Activity and adaptation start at zero before stationary warmup.

**Table 1:** Population parameters. Time constants are in seconds; cell counts and adaptation strengths are dimensionless. Recurrent and inhibition gains use the model normalization.

| Parameter | Symbol | DC | MEC grid | HPC place |
| --- | --- | --- | --- | --- |
| Cells per module | $N$ | 128 | $20 \times 20$ | $80 \times 80$ |
| Activity time constant (s) | $\tau$ | 0.01 | 0.01 | 0.001 |
| Adaptation time constant (s) | $\tau_v$ | 0.1 | 0.1 | 0.002 |
| Recurrent strength | $J_0$ or $J_k$ | 8.75 | 5 | 2, 4 |
| Direct-sum inhibition | $k_h, k_g, \kappa_p$ | 0.05 | 1 | $1 \times 10^{-4}$ |
| Adaptation strength | $m$ | 0.3–0.8 | 0.5 | 0.5 |

#### Discrete updates

All neural populations, including the band cells, advance by forward Euler steps of Δ*t* = 0.001 s. Within each population, activity and adaptation increments use the preceding states; updated synaptic activity is clipped at zero before firing rates are recomputed. Each step updates DC activity, then the band-cell reference and MEC activity, and finally HPC activity. The resulting HPC output and the sensory observation at the current physical position supply that step’s synaptic-learning update.

#### Theta modulation of anchoring inputs

DC and MEC populations share a theta clock with a 0.1 s period (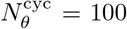 neural steps). Its phase *ϕ* advances uniformly, and the modulation strength *ϑ* controls the anchoring gain:

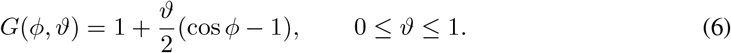

The DC and MEC modulation strengths are *ϑ*_*h*_ and *ϑ*_*g*_, respectively. Zero modulation leaves the anchor constant, whereas full modulation lets it vanish at the weakest phase. Each cycle starts at that phase without resetting neural activity.

#### DC: anchoring activity to heading

Following Ji et al. (2025), a DC ring represents the direction that guides each MEC sweep. The ring contains *N*_*h*_ cells with uniformly spaced preferred directions *α*_*a*_. Cells with nearby preferred directions excite one another through

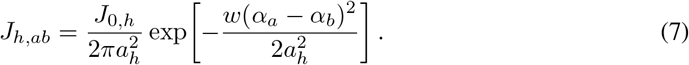

Here *a* and *b* index cells in the ring, *J*_0,*h*_ sets recurrent strength, and *a*_*h*_ = 0.4 rad is the angular width. The wrapping function *w*(*q*) = ((*q* + *π*) mod 2*π*) − *π* maps phase differences to [−*π, π*) rad and acts componentwise on vectors. Recurrence is a direct sum over DC firing rates.

A heading-centered input anchors the bump to the animal’s direction:

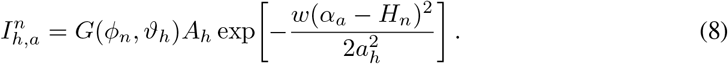

The heading cue is *H*_*n*_ at step *n*, and its amplitude is *A*_*h*_ = 90 a.u. The input and recurrent kernel have the same angular width. As the anchor weakens over theta phase *ϕ*_*n*_, adaptation allows the bump to move away from the heading cue. Module-dependent adaptation strengths *m*_*h,k*_ are listed in Table 2 and implement the scale-dependent sweep-angle organization considered by Ji et al. (2026).

#### MEC: anchoring activity to position

Band-cell integration supplies a reference phase vector *ψ*_*m*_ to MEC module *m* (see Appendix A.4 for the integration mechanism). Its firing-rate vector is *r*_*g,m*_. For grid cell *a* with preferred phase *θ*_*a*_, the reference produces the anchoring field

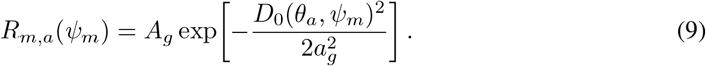

The reference has amplitude *A*_*g*_ = 6 a.u. and phase width *a*_*g*_ = 1 rad. The function *D*_0_ measures distance on the oblique phase sheet. For phase vectors *θ* and *ψ, D*_0_(*θ, ψ*)^2^ = *δ*^*⊤*^*Qδ*, with *δ* = *w*(*θ* − *ψ*). The band-axis metric is

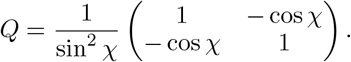

The angle *χ* = *π/*3 rad separates the band axes to match the hexagonal lattice geometry of grid-cell firing fields; ⊤ denotes transpose. Within the periodic MEC sheet, the kernel uses componentwise wrapped differences without searching neighboring images.

MEC recurrent excitation couples cells with nearby preferred phases:

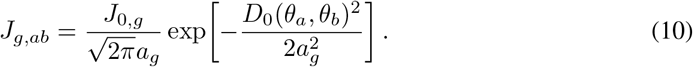

The gain *J*_0,*g*_ sets recurrent strength. Recurrence is a periodic sum over grid-cell firing rates, using the same phase width as the reference field.

A conjunctive direction–position pathway adds shifted copies of the grid activity, weighted by DC firing rates. After the DC update, the MEC input is

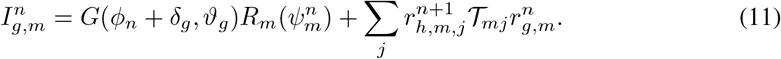

Here *R*_*m*_ is the vector of reference inputs, 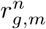 is the preceding grid-rate vector, and 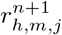 is the updated rate of DC cell *j* in module *m*. The offset *δ*_*g*_ = −4*π/*9 rad shifts the grid anchor relative to the common theta phase. The Fourier operator *T*_*mj*_ shifts the grid pattern by (0.19 rad) 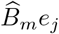, where *e*_*j*_ = (cos *α*_*j*_, sin *α*_*j*_)^*⊤*^ is the preferred DC direction and 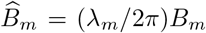. Here *λ*_*m*_ is grid spacing, and *B*_*m*_ maps physical displacement to band-phase displacement; see Appendix A.2 for their definitions and spatial settings. This input steers the grid bump away from its position reference as the theta anchor weakens.

#### HPC: following MEC sampling activity

HPC populations follow the input from MEC without a separate position anchor or theta gate. Their shorter activity and adaptation time constants (Table 1) are chosen to support rapid following of the changing MEC sampling pattern.

CA3 recurrence uses the Gaussian connection kernel in Equation 1, with strength *J*_*k*_ and width *a*_*p,k*_ shared within population *k*. On the 80 × 80 field-center lattice in normalized coordinates [0, 1]^2^, the discrete recurrent input is

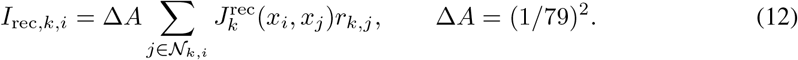

The square neighborhood *N*_*k,i*_ extends ⌈4*a*_*p,k*_/*h*⌉ lattice steps along each coordinate, where *h* = 1*/*79 and *a*_*p,k*_ is expressed in model length units. Rates outside the arena are zero; zero-padded FFT convolution implements this finite, nonperiodic input. Inhibition pools rates within each HPC population, and HPC output drives sensory learning without readout feedback.

### A.2 Spatial scales and grid-to-place projections

HPC field centers uniformly sample the arena [0 m, 10 m]^2^. The spatial computations in this subsection use normalized coordinates in [0, 1]^2^, with one model length unit equal to 10 m: a normalized coordinate or displacement *q* is displayed in physical units as *q*_m_ = (10 m)*q*. MEC preferred phases sample [−*π* rad, *π* rad)^2^.

The matrix *P*_*m*_ maps the firing-rate vector *r*_*g,m*_ of MEC module *m* to the HPC sheet. HPC population *k* receives a dominant input from its paired module and weaker inputs from the remaining modules:

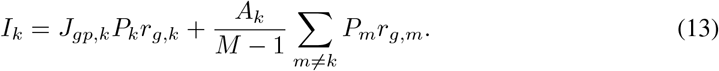

The dominant-input gain is *J*_*gp,k*_, and the auxiliary gain *A*_*k*_ is divided equally among other sources. The common HPC field centers make *P*_*m*_ independent of the target population.

#### From physical position to grid phase

The grid phase corresponding to an HPC field center *x* is

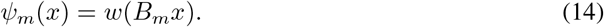

For grid spacing *λ*_*m*_, lattice orientation *γ*_*m*_, and band-axis separation *χ*, the position-to-phase matrix is

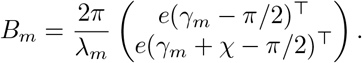

Here *e*(*α*) = (cos *α,* sin *α*)^*⊤*^. Each row projects position along a band direction and converts distance to phase.

#### From phase distance to connection strength

For HPC cell *i* at *x*_*i*_ and MEC cell *a* with preferred phase *θ*_*a*_, the wrapped phase difference is Δ_*m,ia*_ = *w*(*ψ*_*m*_(*x*_*i*_) − *θ*_*a*_). The projection onto the nonperiodic HPC sheet uses the nearest periodic image of the source MEC phase:

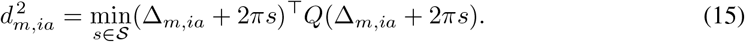

Each integer vector *s* ∈ *S* = {−1, 0, 1} ^2^ selects the central MEC phase image or one of its eight neighbors, without wrapping HPC field centers across arena boundaries. The shortest distance *d*_*m,ia*_ sets the Gaussian connection weight:

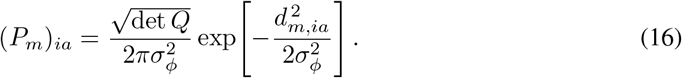

All modules use projection phase width *σ*_*ϕ*_ = 1 rad. The determinant det *Q* sets the Gaussian amplitude in oblique coordinates.

A MEC bump produces several HPC input peaks because grid phases are periodic. Different module spacings help distinguish these locations, and CA3 recurrence helps stabilize a localized bump as the input moves.

#### Spatial scales and input gains

Place-cell field-size diversity motivates multiple HPC scales (Kjelstrup et al., 2008). The relation *a*_*p,k*_ = *λ*_*k*_/(2*π*) and recurrent gains are chosen to let localized HPC bumps follow their dominant MEC inputs at matched scales. This pairing, width relation, and gain choice are modeling assumptions. Table 2 lists the scale-dependent settings.

**Table 2:** Module parameters in increasing grid spacing. One model length unit equals 10 m. Physical lengths and angles carry units in the headers; adaptation is dimensionless and connection gains use model normalization.

| Module | $\lambda_k$ (model) | $\lambda_k$ (m) | $\gamma_k$ (rad) | $a_{p,k}$ (m) | $m_{h,k}$ | $J_k$ | $J_{gp,k}$ | $A_k$ |
| --- | --- | --- | --- | --- | --- | --- | --- | --- |
| M1 | 0.12 | 1.2 | 0 | 0.191 | 0.3 | 2 | 264.873 | 90 |
| M2 | 0.16 | 1.6 | $\pi/30$ | 0.255 | 0.425 | 2 | 198.655 | 60 |
| M3 | 0.2 | 2 | $\pi/15$ | 0.318 | 0.55 | 2 | 158.924 | 60 |
| M4 | 0.24 | 2.4 | $\pi/10$ | 0.382 | 0.675 | 2 | 132.437 | 60 |
| M5 | 0.28 | 2.8 | $2\pi/15$ | 0.446 | 0.8 | 4 | 113.517 | 60 |

### A.3 Sensory encoding and synaptic learning

#### Sensory encoding

The sensory targets encode object-relative vectors or distances to the arena boundary. All three encodings contain 2304 channels with spatial tuning width *σ* = 0.6 m. Positions and distances in this subsection are expressed in metres. The four objects occupy the combinations of 2.5 m and 7.5 m along the two coordinate axes. Table 3 compares the preferred-vector sampling and target amplitudes.

**Table 3:**
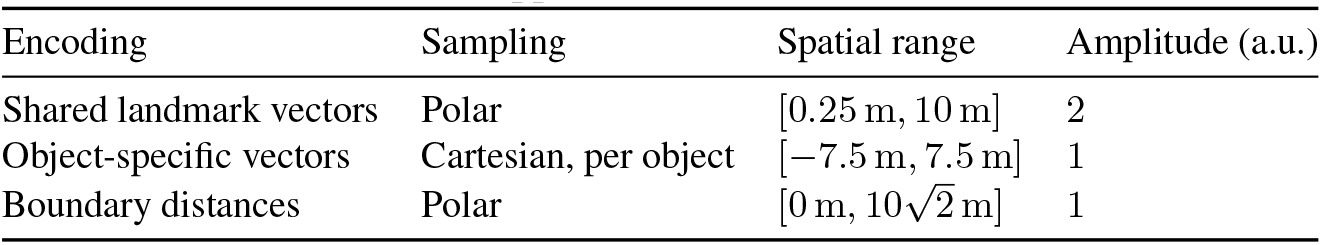
Sensory-encoding parameters. All spatial ranges are in metres; amplitudes are in arbitrary model units (a.u.). The Cartesian range applies to each coordinate.

| Encoding | Sampling | Spatial range | Amplitude (a.u.) |
| --- | --- | --- | --- |
| Shared landmark vectors | Polar | [0.25 m, 10 m] | 2 |
| Object-specific vectors | Cartesian, per object | [-7.5 m, 7.5 m] | 1 |
| Boundary distances | Polar | [0 m, $10\sqrt{2}$ m] | 1 |

For an object at *l*_*i*_, a channel with preferred displacement *ρ* responds at position *x* as

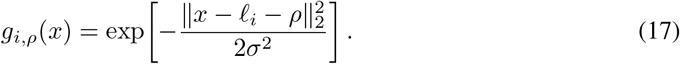

Here *i* indexes objects. Shared landmark-vector cells pool over object identity. Their preferred vectors are *ρ*_*hj*_ = *d*_*h*_(cos *θ*_*j*_, sin *θ*_*j*_), where *h* indexes distance and *j* indexes direction. The shared landmark-vector and boundary encodings each use 36 uniformly spaced preferred distances, including both endpoints of the ranges in Table 3, and 64 uniformly spaced preferred directions around the circle. The shared target is 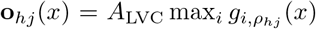, where *A*_LVC_ sets its amplitude. Every object contributes, with no occlusion rule.

Object-specific vectors preserve identity by assigning each object a separate block of channels, with target **o**_*i,ρ*_(*x*) = *g*_*i,ρ*_(*x*). Each object has a 24 × 24 grid of preferred displacements *ρ*, sampled uniformly over the Cartesian range in Table 3, including its endpoints. The four object blocks are concatenated to form the target.

Boundary-distance cells use the same preferred directions. For the unit direction *e*_*θ*_ = (cos *θ,* sin *θ*), *d*(*x, θ*) is the first nonnegative distance from *x* to the square boundary along that ray. It is the smallest valid componentwise intersection distance: (*L* − *x*_*c*_)*/e*_*θ,c*_ when *e*_*θ,c*_ *>* 0, or − *x*_*c*_/*e*_*θ,c*_ when *e*_*θ,c*_ *<* 0. Here *L* = 10 m is the arena side length and *c* indexes Cartesian coordinates; direction components with magnitude below 10^−7^ are omitted. With preferred distance *d*_*h*_, the target is

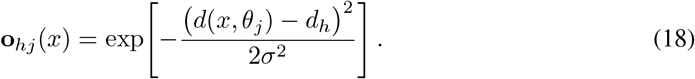

Encoding panels display activity relative to each encoding’s amplitude; training uses the original amplitudes.

#### HPC output and synaptic learning

The mapping *z*_*t*_ = *f* (*r*_*t*_) converts HPC firing rates into the activity presented to the sensory readout. Within population *k*, cell *i* has rate *r*_*k,i*_. Its population has total activity *S*_*k*_ =∑ _*i*_ *r*_*k,i*_ and peak activity *p*_*k*_ = max_*i*_ *r*_*k,i*_. Suppressing the time index, define the peak-normalized rate *r*_*k,i*_ = *r*_*k,i*_/ max(*p*_*k*_, *ϵ*_feat_). The output is

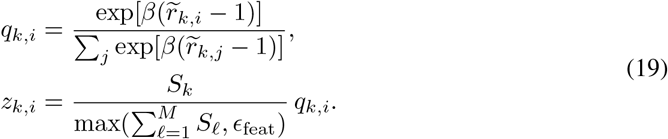

Here *q*_*k,i*_ is the within-population output distribution. The nonlinear transformation controls how many cells contribute appreciably to the readout: small *β* spreads output across many cells, whereas large *β* concentrates it on a few. We use *β* = 8 as an intermediate choice. The numerical floor *ϵ*_feat_ = 10^−30^ prevents division by zero. The indices *j* and *l* run over cells and populations, respectively. Weighting by *S*_*k*_ preserves relative population activity; the concatenated output has unit mass when total HPC activity is at least the numerical floor and vanishes when the circuit is silent.

For the default five HPC populations, concatenation gives 32000 outputs, and the readout matrix has dimensions 2304 × 32000. This matrix starts from zero weights and is the only plastic connection during exploration. Weights may be positive or negative. Each observation produces a prediction and a delta-Hebbian update using the same *z*_*t*_, without theta gating. Equation 4 uses

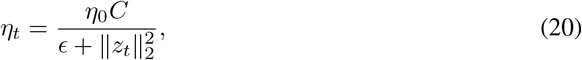

where *η*_0_ is the coefficient selected for each condition and *ϵ* = 10^−8^ prevents division by zero. The reference energy *C* = 1.580 × 10^−2^ was calibrated on a shared-DC trajectory with three MEC modules and then held fixed across experiments, including changes in sensory width and readout population count. See Appendix B.1 for the selected coefficients and selection procedure.

### A.4 Band-cell path integration

#### Band-cell integration and the grid reference

Each MEC module receives a position reference from two band-cell rings, which integrate velocity along its band axes. Their decoded phases form the vector *ψ*_*m*_ that anchors the grid sheet. This construction draws on the band-cell path-integration mechanism in Section 3 and Appendix F.1 of Chu et al. (2025). Periodic phase coding of displacement is also used in hybrid oscillatory-interference and continuous-attractor models (Bush & Burgess, 2014).

#### Neural band integrator

Each band ring contains *N*_*b*_ = 180 cells with uniformly spaced preferred phases *α*_*a*_. The recurrent weight from cell *c* to cell *a* is

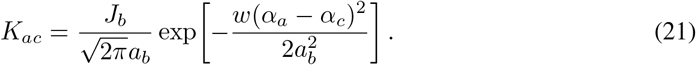

The phase width is *a*_*b*_ = 2*π/*9 rad, and *w* returns the wrapped angular difference. The subscript *b* identifies band-cell quantities; *a* and *c* index the receiving and sending cells. With ring density *ρ*_*b*_ = *N*_*b*_/ (2*π*) cells per radian and inhibition coefficient *k*_*b*_ = 5 × 10^−4^, recurrent strength is set to 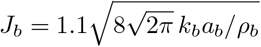.

Velocity ẋ, expressed in model length units per second, produces phase velocity *ν* = (*B*_*m*_ẋ)_*j*_ in rad/s along axis *j* of module *m*, where *B*_*m*_ is the position-to-phase matrix. Positive and negative velocities recruit opposite shifts of the recurrent input:

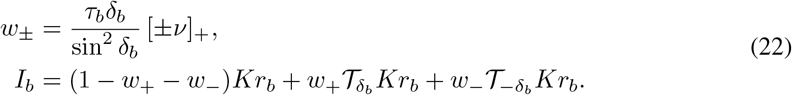

The vector *r*_*b*_ contains band-cell firing rates, *I*_*b*_ is their recurrent input, and *τ*_*b*_ = 0.001 s is the activity time constant. The operator *T*_*δ*_ translates a ring pattern by phase *δ* through a Fourier shift. The fixed shift *δ*_*b*_ = 0.05 rad sets the displacement of each connection pattern, while *w*_+_ and *w*_*−*_ determine its velocity-dependent contribution. Rectification is [*v*]_+_ = max(*v,* 0).

Band cells have no adaptation. Their synaptic activity and firing rates follow

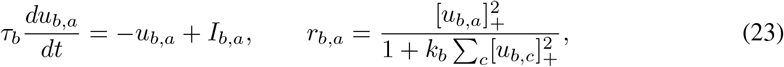

with inhibition pooled within each ring. Synaptic activity begins as 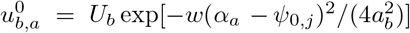, where *U*_*b*_ = 30 a.u. and *ψ*_0,*j*_ is the starting phase along axis *j*. A 2 s stationary warmup allows the bump to settle.

The decoded phase is arg 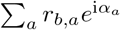, where i is the imaginary unit and arg returns the complex argument. These phases are unwrapped through time and converted to displacement using 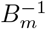. Decoding reads the neural state without reconstructing it. Initial alignment fixes the position origin; subsequent physical positions do not correct the integrated estimate. Band states and constants use double precision.

## B Experimental environment, training, and evaluation

### B.1 Experimental conditions and training

Conditions are paired by physical trajectory and sensory observations. Positions, times, and angles use metres, seconds, and radians; the arena is [0 m, 10 m]^2^. Unless varied below, circuit settings, sensory targets, and synaptic learning follow Appendix A, with full theta modulation and the selected sweep-model learning coefficient.

#### Learning conditions

Figure 3 compares the full model, a no-sweep control, and Vector-HaSH. The no-sweep control uses zero theta modulation, *ϑ*_*h*_ = *ϑ*_*g*_ = 0. Its separately selected learning coefficient is *η*_0_ ≃ 0.149; the sweep model uses 0.53836008. The theta-strength scan varies the anchoring modulation and uses the sweep-model learning coefficient at every strength, including zero. Vector-HaSH matches population sizes, training observations, and evaluation queries; see Appendix B.4 for its representation and readout.

#### Training and trajectories

Readout training uses every neural time step after 2 s of stationary warmup, using the synaptic learning rule in Equation 4. Table 4 lists durations and seeds; Appendix B.2 defines the trajectories. Validation averages errors over seeds 142–144; independent confirmation trains from zero with seeds 242–244 and no further parameter selection.

**Table 4:** Simulation conditions. Durations exclude warmup; each seed identifies one independent trajectory.

| Figure | Duration; seeds | Condition or measurement |
| --- | --- | --- |
| 2c | 400 s; 42–44 | Band integration; no readout learning |
| 2d | 3 s; prescribed | One DC–MEC pair; prescribed motion |
| 2e | 4 s; 42 | Five modules; no readout learning |
| 3 | 400 s; 42–44 | Sweeps, no sweeps, Vector-HaSH |
| 4 | 400 s; 42 | HPC M1 versus all populations; recurrence $J = 0$ |
| 5; 7d | 4,000 s; 42 | Fixed zero; full theta; descending schedule |
| 6a | 400 s; 42 | $\vartheta \in \{0, 0.25, 0.5, 0.75, 1\}$ |
| 6b–d | 400 s; 142–144 | Adaptation × sensory width; first 20 s for span |
| 7a | 400 s; 142–144 | Separate coefficient searches |
| 7b,c,i | 400 s; 42–44 | Regional errors (b,c); encoding comparisons (i) |
| 7e–h | One position | Sensory definitions in Appendix A.3 |
| 8 | 2 s (20 cycles); 42–44 | Two equal grid inputs; $J = 0, 2$; no learning |

#### Learning-coefficient selection

For the sweep and no-sweep conditions in Figure 3, selection minimizes mean all-position error at 400 s. The coarse candidates are 0.025, 0.05, 0.1, 0.2, 0.4, 0.8, 1, 1.6, 3.2. Each search adds *N*_*f*_ = 6 coefficients between the neighbors of its best coarse candidate: [0.2, 0.8] with sweeps and [0.1, 0.4] without sweeps. For bracket endpoints *a, b*,

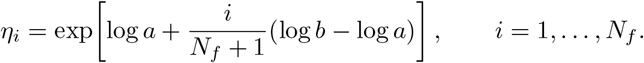

Fine candidates are rounded to eight decimal places. The best of all fifteen candidates sets the condition-specific coefficients given above, which are held fixed thereafter.

#### Sensory-encoding comparison

The shared-LVC, object-specific-vector, and boundary-distance conditions share HPC activity, output count, and learning coefficient; only the sensory targets differ.

### B.2 Exploration trajectories and activity examples

A waypoint controller generates exploration independently of neural activity and prediction error, favoring underexplored locations and slowing near walls and turns. NumPy’s `default_rng(seed)` sets initial conditions, modulation phases, and waypoints. Initial position is uniform on [3 m, 7 m]^2^, and heading is uniform on [−*π, π*] rad, with speed 0.28 m/s and zero angular velocity. The first target is uniform on [1.5 m, 8.5 m]^2^; later targets occupy a 12 × 12 lattice on [0.8 m, 9.2 m]^2^. On reaching or expiring a target, the highest-scoring eligible waypoint is selected using

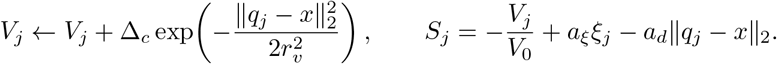

Here *q*_*j*_ is a waypoint, *x* the current position, *V*_*j*_ its proximity-weighted visit time, Δ_*c*_ the controller interval, and *ξ*_*j*_ an independent standard Gumbel draw. Table 5 gives the spatial scales, score coefficients, and target criteria.

**Table 5:**
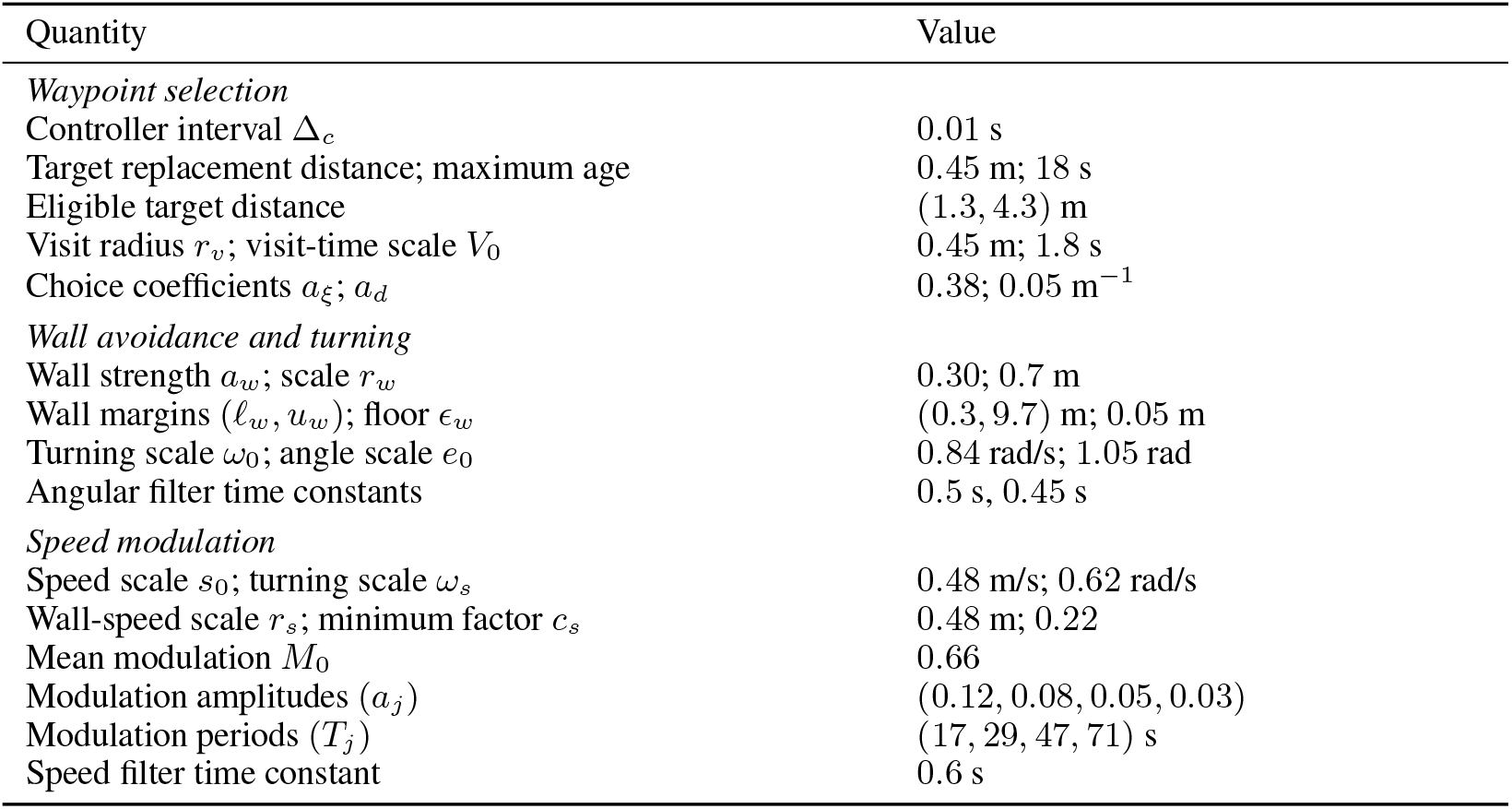
Exploration-controller parameters in physical units. Units are given for all dimensional quantities; values without units are dimensionless.

| Quantity | Value |
| --- | --- |
| *Waypoint selection* |  |
| Controller interval $\Delta_c$ | 0.01 s |
| Target replacement distance; maximum age | 0.45 m; 18 s |
| Eligible target distance | (1.3, 4.3) m |
| Visit radius $r_v$; visit-time scale $V_0$ | 0.45 m; 1.8 s |
| Choice coefficients $a_\xi$; $a_d$ | 0.38; 0.05 m<sup>-1</sup> |
| *Wall avoidance and turning* |  |
| Wall strength $a_w$; scale $r_w$ | 0.30; 0.7 m |
| Wall margins $(\ell_w, u_w)$; floor $\epsilon_w$ | (0.3, 9.7) m; 0.05 m |
| Turning scale $\omega_0$; angle scale $e_0$ | 0.84 rad/s; 1.05 rad |
| Angular filter time constants | 0.5 s, 0.45 s |
| *Speed modulation* |  |
| Speed scale $s_0$; turning scale $\omega_s$ | 0.48 m/s; 0.62 rad/s |
| Wall-speed scale $r_s$; minimum factor $c_s$ | 0.48 m; 0.22 |
| Mean modulation $M_0$ | 0.66 |
| Modulation amplitudes $(a_j)$ | (0.12, 0.08, 0.05, 0.03) |
| Modulation periods $(T_j)$ | (17, 29, 47, 71) s |
| Speed filter time constant | 0.6 s |

To steer away from walls, a repulsive vector is added to the unit direction toward the target:

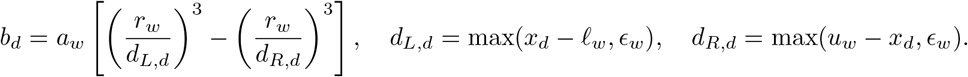

The wall margins *l*_*w*_, *u*_*w*_ and floor *ϵ*_*w*_ bound the repulsion. The resulting direction gives the wrapped heading error *e*. Two successive first-order filters transform *ω*_*\**_ = *ω*_0_ tanh(*e/e*_0_) into angular velocity *ω*. Desired speed is

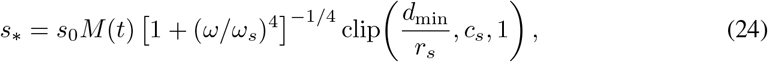

where *d*_min_ is the nearest wall distance and clip(*v, a, b*) = min(*b,* max(*a, v*)). Speed modulation is *M* (*t*) = *M*_0_ + ∑_*j*_ *a*_*j*_ sin(2*πt/T*_*j*_ + *φ*_*j*_), with independent *φ*_*j*_ *~ U* [0, 2*π*). A final first-order filter smooths speed; position advances along the midpoint heading.

Cubic splines interpolate positions to neural sampling times; tangents and position differences give heading and velocity. Accepted trajectories stay inside the arena, below 0.5 m/s and 1 rad/s.

#### Activity examples

Figure 2c tests the band integrator in Appendix A.4. Panel d uses one DC– MEC pair with *m*_*h*_ = 0.6, grid spacing 2 m, and orientation 0 rad; motion starts at (0.8, 0.9) m at 0.4 m/s. Panel e shows two theta cycles of the five-population circuit in animal-relative coordinates rotated to align heading upward.

### B.3 Stationary evaluation and spatial measurements

#### Checkpoint queries

Learning is disabled for evaluation on a 51 × 51 query grid *G* including the arena boundaries. Each query resets MEC/HPC activity and adaptation, anchors the circuit to the query location, and settles for 2.8 s with zero velocity, heading, and theta modulation. Stationary queries use only the position anchor as external MEC input. Adaptation parameters retain their training values. The final HPC rates give *z* = *f* (*r*) (Equation 19) for the stored readout; Vector-HaSH uses its fixed features. Sensory targets enter only the error calculation.

#### Prediction error and replication

Relative error at query *y* and its mean over query set *S* at checkpoint *T* are

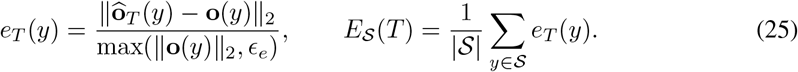

Here **ô**_*T*_ is the prediction and *ϵ*_*e*_ = 10^−12^. Trajectories are the independent replicates: each contributes one spatially averaged error per checkpoint. Across *n* trajectories, SD uses denominator *n* − 1 and SEM is 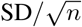. Learning curves connect checkpoints without smoothing.

#### Visited and unvisited regions

Visited and unvisited positions refer to the complete trajectory, so a visited position may be reached after the evaluation checkpoint. For total duration *T*_*\**_, the fixed query regions are

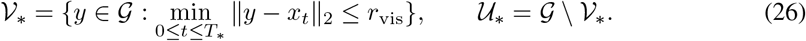

The visit radius is *r*_vis_ = 0.15 m, and all recorded positions, including the start, contribute. The same regions are used at every checkpoint and for all paired conditions, so regional curves compare the same locations throughout learning. Regional errors combine as

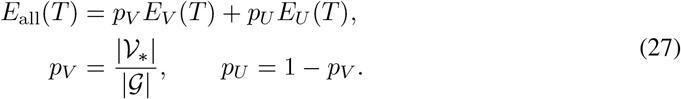

Here *E*_*V*_, *E*_*U*_ are regional errors and *p*_*V*_, *p*_*U*_ are constant query-grid fractions for each trajectory. Weighting precedes averaging across trajectories.

#### Physical and HPC coverage

Coverage uses a 100 × 100 lattice of arena cell centers and radius *r*_vis_. Physical coverage accumulates positions from the start through checkpoint *T*; HPC coverage accumulates the maximum-rate cell position at each sample, over M1 or the union of M1–M5. Peaks at or below *ϵ*_*r*_ = 10^−12^ are excluded. This cell-center lattice differs from the boundary-inclusive error-query grid.

#### HPC sweep span

For the width scans, the maximum-rate cell in population *k* gives position *x*_*k,t*_ and animal-relative displacement 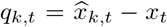. A decrease greater than *π* rad in wrapped grid theta phase marks a cycle boundary. Only complete 0.1 s cycles are retained. Span within cycle *c* is

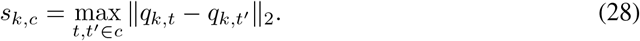

Valid samples have finite rates, with peak and total activity above *ϵ*_*r*_; a cycle requires at least two valid samples. Spans are averaged over cycles within each population and trajectory, then across trajectories.

### B.4 Vector-HaSH baseline and matched comparison

We adapt the Vector-HaSH grid-to-HPC representation and pseudoinverse sensory association of Chandra et al. (2025) to the continuous arena, matching the five grid spacings and the numbers of grid, HPC, and sensory neurons. The baseline uses square, periodic, axis-aligned grid modules with spacings (1.2, 1.6, 2.0, 2.4, 2.8) m. Each module has *N*_*g*_ = 20 cells per axis, with dimensionless preferred phases *p*_*a*_ ∈ [0, 1)^2^ and response

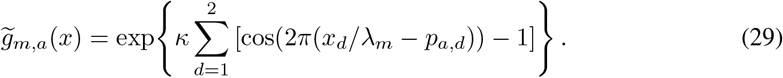

Here *λ*_*m*_ is module spacing, *a* indexes cells, *d* spatial coordinates, and *κ* controls tuning concentration. Normalized modules 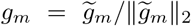 are concatenated into *g*. HPC activity is *h*(*x*) = [*W*_*pg*_*g*(*x*) − *b*_*h*_]_+_, with a fixed 32000 × 2000 projection *W*_*pg*_ drawn from *N* (0, 1) using the trajectory seed, threshold *b*_*h*_ = 2.5, and componentwise rectification [*v*]_+_ = max(*v*, 0). HPC activity has no further normalization or recurrent refinement.

Grid concentration is selected by mean all-position error at 400 s over the validation trajectories in Appendix B.1. Coarse candidates are *κ* ∈ {0.5, 1, 2, 4, 8}, followed by geometric midpoints 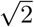 and 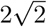 around the best coarse value, 2. The selected 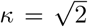 is fixed for subsequent comparisons. Population sizes, spacings, threshold *b*_*h*_, and the random projection within each seed remain fixed during this search.

The initially zero readout is fitted at each checkpoint to all preceding observations, including repeated visits. HPC activities and sensory targets form rows of *X* ∈ ℝ^*n×d*^ and *Y* ∈ ℝ^*n×p*^, where *d, p* are population sizes. The least-squares map is 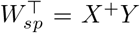; the stored *W*_*sp*_ ∈ ℝ^*p×d*^ acts on column activity vectors. Computation uses the eigendecomposition of *X*^*⊤*^*X* in float64, retaining eigenvalues strictly greater than 10^−12^ times the largest eigenvalue. Equivalently, singular values of *X* at or below 10^−6^ times its largest singular value are discarded. Retained modes are inverted without ridge regularization.

### B.5 HPC population number and CA3 recurrent-excitation ablations

Figure 4a,b uses M1’s 6400 cells alone for output normalization (Equation 19) and sensory readout, compared with 32000 cells across all populations. M1 retains its dominant and auxiliary grid inputs (Table 2). Coverage follows Appendix B.3.

Panels c,d set all CA3 recurrent strengths *J*_*k*_ to zero, retaining inhibition, adaptation, time constants, and grid inputs. Panel d shows M1, M3, and M5 at 10.08 s after warmup (theta phase 0.489 rad). Snapshot rows use separate color scales.

#### Two-module continuity test

Figure 8 supplies two HPC populations with equal inputs from grid modules M1 and M2. Their original gains to HPC M1 are summed and divided equally, giving 143.687 per input (display-rounded). Grid rates are not normalized separately. The populations use M1 geometry and differ only in recurrent strength, *J* = 0 or 2, with zero initial activity and 2 s stationary warmup. There is no sensory learning.

A non-local jump is a displacement exceeding 1 m between consecutive maximum-rate decoded positions within a theta cycle. Cycle-boundary transitions are counted separately.

### B.6 Theta-annealing protocol

Figure 5 compares fixed theta strengths *ϑ* = 0 and 1 with a prescribed descending schedule over the same 4,000 s exploration trajectory. All three conditions use the same fixed learning coefficient *η*_0_ = 0.53836008 to compare theta modulation without condition-specific learning-rate tuning. The no-sweep condition corresponds to *ϑ*_*h*_ = *ϑ*_*g*_ = 0.

The schedule sets *ϑ*_*h*_ = *ϑ*_*g*_ to 1, 0.8, 0.6, 0.4, 0.2 at 0, 600, 1000, 1400, 1800 s, respectively, holding the last value through 4,000 s. Changes act on the DC and MEC anchor gates (Equation 6) and preserve neural activity, adaptation, theta phase, and readout weights. The schedule depends only on elapsed time.

### B.7 Theta strength, grid adaptation, and sensory-width scans

Figure 6a varies the common DC/MEC theta strength over the values in Table 4, with the conditions defined in Appendix B.1. For this scan and the descending schedule, neural computations outside the band integrator and readout computations use float32.

Panel b crosses MEC adaptation 0.2–0.6 in steps of 0.025 with shared-LVC widths *σ* ∈ {0.2, 0.3, 0.4, 0.5, 0.6} m. Adaptation changes identically in all MEC modules; DC/HPC adaptation stays at default. Widths share one neural sequence per adaptation value and seed but use separate targets and readouts. The learning coefficient and reference energy are fixed across widths.

For each width, panel c selects the adaptation minimizing mean final all-position error and reports its sweep span averaged across the five HPC populations. Panel d retains every adaptation value and reports population-specific spans. Span measurement and averaging follow Appendix B.3.

## C Additional experimental results

### C.1 Learning-coefficient selection and independent confirmation

Separate learning-coefficient searches give different optima with and without sweeps (Figure 7a). With these coefficients fixed, independent-confirmation errors are 0.511 ± 0.002 with sweeps versus 0.620 ± 0.002 without (mean ± SEM). The sweep advantage therefore persists after tuning and on trajectories excluded from selection.

### C.2 Regional learning and sensory-encoding comparisons

Figure 7b,c compares learning within and outside the region covered by each complete 400 s trajectory. The regions remain fixed across checkpoints, so the visited set includes positions reached later in exploration; see Appendix B.3 for the region definitions. Vector-HaSH reaches the lowest final error at visited positions, while its unvisited error initially rises before declining. The sweep model has the lowest unvisited error.

Figure 7d extends the regional comparison to 4,000 s, using locations that remain unvisited over this longer trajectory. During later exploration, full theta modulation (*ϑ* = 1) gives lower error at these locations than either zero modulation or the descending schedule, while the schedule reaches lower all-position error (Figure 5c). Because all-position error is a weighted mean of visited and unvisited errors (Equation 27), the schedule’s overall advantage over full modulation reflects greater accuracy in the visited region.

To test whether the learning benefit depends on the sensory representation, Figure 7e–h shows three encodings at the same position in one scene. Panel e locates the animal and objects; panels f–h show responses that pool object-relative vectors across objects, preserve object identity in separate channel groups, or encode distances to the boundary. With the neural activity and learning coefficient matched across encodings (Appendix B.1), sweeps reduce all-position prediction error for all three targets (panel i).

### C.3 Recurrence and spatial continuity with two grid modules

Two periodic grid inputs can support competing locations. Their strongest overlap can remain nearby or switch to a distant location as the activity peaks move (Figure 8a). Local recurrent excitation favors CA3 cells near the preceding activity peak, helping maintain the same local position representation when the grid input is ambiguous.

Without recurrence, the decoded peak switches between distant locations; with recurrence, it moves locally under the same inputs (Figure 8b,c). Recurrence eliminates within-cycle jumps across all three trajectories (panels d,e), reducing the maximum step from 5.10 m to 0.179 m. Neither condition has cycle-boundary jumps.

